# Verbal working memory and syntactic comprehension segregate into the dorsal and ventral streams

**DOI:** 10.1101/2024.05.05.592577

**Authors:** William Matchin, Zeinab K. Mollasaraei, Leonardo Bonilha, Chris Rorden, Gregory Hickok, Dirk den Ouden, Julius Fridriksson

**Affiliations:** Dept. of Communication Sciences and Disorders, University of South Carolina; Dept. of Pharmacology, Physiology, Neuroscience, University of South Carolina; Dept. of Psychology, University of South Carolina; Dept. of Cognitive Sciences, Dept. of Language Science, University of California, Irvine

**Author notes:** Corresponding author 915 Greene St., Discovery 1, Room 202D, Columbia, SC 29208, USA.

**Keywords:** aphasia, lesion-symptom mapping, tractography, working memory, syntax, rhyme judgment, sentence comprehension

## Abstract

Syntactic processing and verbal working memory are both essential components to sentence comprehension. Nonetheless, the separability of these systems in the brain remains unclear. To address this issue, we performed causal-inference analyses based on lesion and connectome network mapping using MRI and behavioral testing in 103 individuals with chronic post-stroke aphasia. We employed a rhyme judgment task with heavy working memory load without articulatory confounds, controlling for the overall ability to match auditory words to pictures and to perform a metalinguistic rhyme judgment, isolating the effect of working memory load. We assessed noncanonical sentence comprehension, isolating syntactic processing by incorporating residual rhyme judgment performance as a covariate for working memory load. Voxel-based lesion analyses and structural connectome-based lesion symptom mapping controlling for total lesion volume were performed, with permutation testing to correct for multiple comparisons (4,000 permutations). We observed that effects of working memory load localized to dorsal stream damage: posterior temporal-parietal lesions and frontal-parietal white matter disconnections. These effects were differentiated from syntactic comprehension deficits, which were primarily associated with ventral stream damage: lesions to temporal lobe and temporal-parietal white matter disconnections, particularly when incorporating the residual measure of working memory load as a covariate. Our results support the conclusion that working memory and syntactic processing are associated with distinct brain networks, largely loading onto dorsal and ventral streams, respectively.

## Introduction

Language contains multiple interlocking relationships among the elements that make up sentences. Syntax, the ability to combine these elements into complex structures, allows for the expression of novel ideas. However, processing these complex structures requires significant processing resources. For example, in noncanonical structures like object-relatives (e.g. *the dog*_*1*_ *that [the cat is chasing* — _*1*_*] is super cute)*, embedded relative clauses increase both the linear difference and degree of structural interference between the head noun and its related position within the embedded clause. The increased processing difficulty and neural resources associated with complex noncanonical structures such as these has accordingly been heavily studied.^1–6^ Working memory (WM), the capacity to store and manipulate information over time, is a central aspect of higher-level cognition ^7,8^ and a strong candidate as part of the explanation for these complexity effects.^4,9–11^ Thus, the separability between syntax and WM remains an enduring question in psychology, cognitive neuroscience, and aphasia research, with implications for the nature of linguistic processing and representations, as well as clinical interventions for sentence-processing deficits.^1,2,4,6,11–18^

In healthy subjects, verbal WM is associated with a distributed network of superior temporal, inferior parietal, and inferior frontal cortices,^19–24^ which overlaps with networks activated for sentence processing in functional magnetic resonance imaging (fMRI) studies.^5,25,26^ However, it is largely unclear if WM and sentence processing depend on the same cortical and subcortical networks, since fMRI studies cannot resolve which structures are necessary for or merely correlated with a given process. Causal inference requires different methods, such as lesion-symptom mapping in people with post-stroke aphasia,^27,28^ which also allows for more direct translation between research and clinical practice. In addition, while syntactic processing has been strongly associated with dorsal stream networks in past research,^29–38^ suggesting a close relationship between verbal WM and syntax,^39^ one recent proposal states that syntactic comprehension critically relies only on the ventral stream,^40^ suggesting a categorical segregation of these systems in the brain.

Sentence comprehension deficits are common in post-stroke aphasia, and tend to be more pronounced and chronic than word-level deficits.^41^ A large body of work has associated syntactic comprehension deficits in nonfluent aphasia to frontal-dorsal stream damage,^37,42–47^ whereas several recent lesion-symptom mapping studies have shown that impaired syntactic comprehension is primarily associated with temporal-parietal lesions.^48–55^ WM deficits often involve damage to inferior frontal, parietal, and temporal lobe lesions, similar to the networks identified in functional neuroimaging research. However, the extent of overlap or dissociation of WM and syntactic deficits is unclear.^12,16,56,57^ For example, deficits in receptive syntactic processing often observed in speakers with nonfluent aphasia and frontal damage might be due in part to impaired verbal WM resources in such patients.^14,17,18,58,59^

In this study, we aim to clarify the lesion correlates, both in terms of structural damage and connectivity, associated with syntactic processing ability and verbal WM load—a crucial factor believed to underlie poor performance across various cognitive domains, notably sentence processing. To achieve this, we employ two distinct rhyme judgment tasks, leveraging them to triangulate our findings. Our focus lies on the Triplets part A of the Temple Assessment of Language and Short-term Memory in Aphasia (TALSA) ^60,61^. In the Triplets task, participants evaluate three auditorily-presented words, each paired with a corresponding picture, to identify which pair rhymes, by selecting the appropriate pictures. This task imposes significant verbal WM load, as it necessitates the simultaneous processing and assessment of three different two-word pairs for rhyme judgment. However, it does not require overt speech articulation ability, unlike commonly used span and repetition tasks.^62–66^ To isolate the impact of this processing load, beyond the basic task execution, we utilize the Auditory Word Rhyme Judgment task (RhymeJudge) from the Psycholinguistic Assessments of Language Processing in Aphasia (PALPA)^67^ as a covariate, assessing participants’ general ability in auditory-based metalinguistic rhyme judgment. Additionally, we incorporate the Auditory Word Recognition (AudWordRec) measure from the Western Aphasia Battery-Revised (WAB-R)^65^ to control for participants’ ability to match auditory word stimuli to visual objects. In a subset of participants, we assessed syntactic processing ability through the use of the Noncanonical sentence comprehension, with a covariate for WM load, operationalized as residual performance on the Triplets task after removing variance associated with RhymeJudge and AudWordRec as detailed above. We also conducted behavioral assessments to explore the extent of shared variance between the rhyme judgment tasks and other measures, alongside comparisons with the WAB-R repetition task—a conventional approach to evaluating verbal WM deficits. By employing these methodologies, our study not only addresses the limitations of previous assessments but also sheds light on the intricate relationship between verbal WM load and cognitive performance across various linguistic tasks. In doing so, we contribute to a deeper understanding of the underlying mechanisms of cognitive deficits observed in individuals with aphasia and related disorders.

## Materials and methods

### Participants & measures

Retrospective analyses were conducted on data from 103 participants in a broader clinical trial (clinicaltrials.gov ID: NCT03416738) conducted at the University of South Carolina (USC) and the Medical University of South Carolina (MUSC), which has overlapped with previous publications from our research group.^51–53,68,69^ All procedures were approved by the Institutional Review Boards for both institutions, and informed consent was obtained prior to participation.

Patient demographic info is presented in Table 1. All participants were at least one year post-stroke. This larger study involved a large set of baseline measures, including the *Western Aphasia Battery-Revised* (WAB-R), a large subset of the *Psycholinguistic Assessments of Language Processing in Aphasia* (PALPA), and a large subset of the *Temple Assessment of Language and Short-term Memory in Aphasia* (TALSA). A subset of 78 participants out of the total 103 of these participants were also administered the *Northwestern Assessment of Verbs and Sentences* (NAVS).

**Table 1.** Participant demographics. WAB-R = Western Aphasia Battery-Revised. AQ = aphasia quotient. SD = standard deviation.

| Demographic Variable | Full verbal WM & WAB-R group (N=103) | Subset with noncanonical sentence comprehension (N=78) |
| --- | --- | --- |
| Sex | 41 female, 62 male | 29 female, 49 male |
| Age at testing (years) | 60.63 years (SD = 10.98 years) | 61.09 years (SD = 10.91 years) |
| Days post stroke at testing | 1697 days (SD = 1638 days) | 1883 days (SD = 1732 days) |
| Lesion volume | 118,462 mm<sup>3</sup> (SD = 91,812 mm<sup>3</sup>) | 124,353 mm<sup>3</sup> (SD = 90,889 mm<sup>3</sup>) |
| Aphasia subtypes by WAB-R | 40 Broca's, 24 Anomic, 16 conduction, 10 Not Aphasic by WAB-R, 7 Global, 4 Wernicke's, 1 Transcortical sensory, 1 Transcortical motor | 34 Broca's, 14 Anomic, 11 conduction, 10 Not Aphasic by WAB-R, 6 Global, 3 Wernicke's, 1 Transcortical sensory |
| WAB-R AQ | 62.25 (SD = 24.94) | 60.94 (SD = 26.03) |
| WAB-R Repetition (percent correct) | 56% (SD = 32%) | 55% (SD = 31%) |
| WAB-R Auditory Word Recognition (AudWordRec) (percent correct) | 83% (SD = 22%) | 82% (SD = 22%) |
| TALSA Triplets score (percent correct) | 66% (SD = 22%) | 66% (SD = 23%) |
| PALPA Auditory word rhyme judgment score (RhymeJudge) (percent correct) | 82 % (SD = 14%) | 81% (SD = 14%) |
| NAVS Noncanonical Sentence comprehension score (percent) | - | 70% (SD = 23%) |
| correct) |  |  |

Here we focus on four specific measures contained in these larger test batteries:

- the **Auditory Word Recognition subtest of the WAB-R (AudWordRec)**, with 60 trials, which involves identifying common household objects given an auditorily presented word, e.g. asking the patient: “point to the book”. Measures simple auditory word comprehension and object recognition.
- the **Auditory Word Rhyme Judgment** subtest of the PALPA (**RhymeJudge**), 30 trials, which requires subjects to indicate whether a pair of presented words rhymes or not. Half of the presented items rhymed (e.g. *king, sing*), and half did not (e.g. *beard, heard*). Measures basic phonological processing.
- the **Triplets** Part A (3 comparisons) of the TALSA (**Triplets**), with 30 trials. An array of three simple objects is presented, and the names are presented auditorily (e.g., *mouse, dice, house*). Participants are asked to identify which of two object names rhyme with each other. The non-matching, distractor object name overlapped phonologically and orthographically with the other items. Measures the ability to handle a heavy verbal WM load.
- the **Noncanonical Sentences** (passives, object WH questions and object-relatives) from the Sentence Comprehension Test of the NAVS (**Noncanonical**), consisting of 15 trials (out of the total 30). This task involves matching an auditorily presented sentence (e.g., *The cat is chased by the dog? Who is the cat chasing? Pete saw the dog who the cat was chasing*) with the correct picture out of two, role-reversed alternatives (e.g., a cat chasing a dog or a dog chasing a cat). Assesses high-level linguistic comprehension, including syntactic processing.

These measures were selected to isolate verbal WM load from perceptual, motor, and broader task demands, as well as to differentiate WM load from syntactic processing. The TALSA and PALPA have been previously validated in people with aphasia; performance on the Triplets task is near ceiling in healthy individuals ^61^. In addition, we included the WAB-R Aphasia Quotient and WAB-R Repetition scores for further evaluation of the two rhyme judgment tasks as well as noncanonical sentence comprehension.

The TALSA Triplets measure assesses the extent to which a participant can successfully manage substantial verbal WM load. In this task, three different pairs of words must be held in memory and evaluated for their rhyming status. However, to effectively perform the task, participants must also be able to process an auditory stimulus, perform the metalinguistic rhyme judgment (incorporating both phonological access and executive function ability), and map this onto a visual object representation. By contrast, the WM demands of the PALPA RhymeJudge test are low, as only a single pair of items must be analyzed, with no delay, and no visual object processing is required. This task has similar metalinguistic demands as the TALSA Triplets task. Therefore, by using the WAB-R AudWordRec as a covariate to control for auditory word object recognition abilities and the PALPA RhymeJudge as a covariate to control for metalinguistic rhyme judgment ability, we can obtain greater precision on WM resources than using the Triplets task on its own.

The NAVS Noncanonical sentence comprehension task measures the extent to which a participant can process linguistic structure and identify the picture which contains the correct thematic relationships as specified by the grammatical structure of the sentence. Given that processing noncanonical sentences is associated with verbal WM resources, we include the residual Triplets performance (as described above) as a covariate to control for the ability to process verbal WM load, thereby providing a purer measure of syntactic processing ability.

### Brain imaging and lesion mapping

High-resolution anatomical images and diffusion-weighted images (DWI) were collected at the University of South Carolina and the Medical University of South Carolina on a 3 T Siemens Prisma scanner with a 12-channel head coil. T1 images were collected using an MP-RAGE sequence, 1 mm isotropic with a 256 x 256 matrix, 9° flip angle, TR 2,250 ms, and either 160 slices with TI 925 ms and TE 4.52 ms, or 192 slices with TI 925 ms and TE 4.15 ms, using parallel imaging (GRAPPA = 2, 80 reference lines). T2 images were collected using a 3D-SPACE sequence with 192 slices (1 mm), TR 2800 ms, TE 402 ms, variable flip angle, 256 x 256 matrix, using parallel imaging (GRAPPA = 2, 120 reference lines). DWI scans were acquired using a 210 × 210 matrix, 43 volumes sampling 36 directions with b = 1000 sec/mm2 (seven volumes b = 0), 90° flip angle, TR 5250 msec, TE 80 msec with parallel imaging (GRAPPA = 2, 80 reference lines). Images were acquired twice; phase encoding polarity was reversed between scans.

A binary map of each participant’s lesion was drawn onto their T2 image to highlight lesion areas, by either LB (a neurologist) or RNN (a cognitive neuroscientist) with consultation with CR (expert on brain imaging and lesion analysis). Both individuals were blind to the behavioral data. Maps were then aligned to each participant’s high-resolution T1 image, and lesions were “healed” in the T1 image using the corresponding brain anatomy from the intact hemisphere using SPM12^70^ and custom scripts. The T1 image and its corresponding lesion map were subsequently warped to the standard Montreal Neurological Institute (MNI) template using SPM12. Each lesion map warped to standard space was then re-binarized using a 50% probability threshold. Lesion overlap maps for the full set of 103 participants and for the subset of 78 participants are shown in Figure 1. As can be seen, coverage (and thus statistical power) was highest in the perisylvian area for both sets of participants.

**Figure 1.**
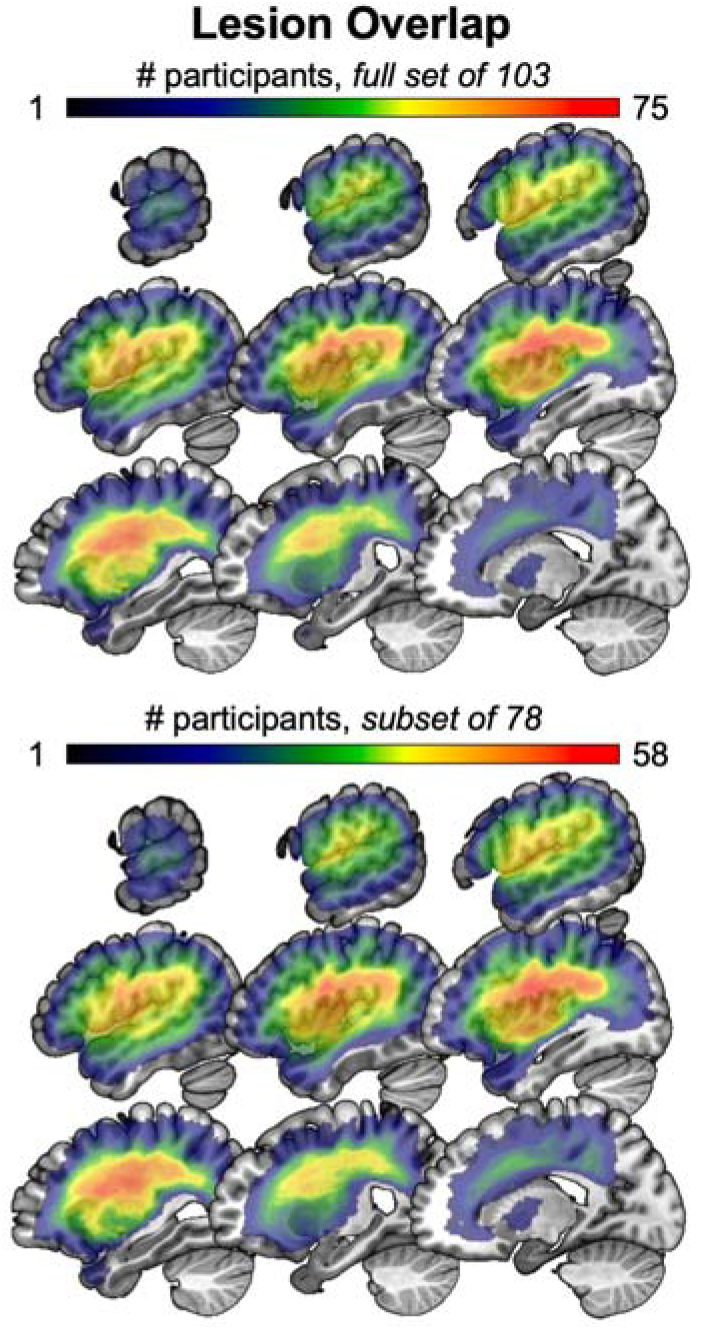
Lesion overlap maps. A: lesion overlap map for the full set of 103 participants who were administered the WAB-R as well as the TALSA and PALPA rhyme judgment tasks. B: lesion overlap map for the subset of 78 participants who also were administered the noncanonical sentence comprehension task.

T1 images were used to align the Johns Hopkins University (JHU) atlas region parcellations ^71^ to DWI images. Then, connection strength between regions was estimated using fiber count, corrected for distance and region volume, for each of the pairwise connections within left hemisphere regions by using probabilistic tractography as implemented in FSL’s Diffusion Toolbox (FDT) method,^72^. Reciprocal connectivity for each pair of regions was calculated as the average of the total fiber count arriving in region A when region B was seeded and vice versa.

The lesioned tissue was removed from all tractography tracings to maximize accuracy of fiber tracking, which also minimizes the effect of lesion volume on the final analyses. The estimated number of streamlines from a fully lesioned region was set to zero (therefore, there is no estimated connectivity from a fully lesioned region). The number of tracked fibers for a pair of regions was divided by the total volume of both regions, which controls for unequal region volumes across connection pairs. This was the final (dis)connection value for each pair of regions in our analyses.

### Analyses

We first performed two-tailed correlation analyses (Pearson’s r) in JASP ^73^ to establish the extent of the relationship among the behavioral variables and assess the extent to which there is distinct variance associated with the rhyme judgment tasks, PALPA RhymeJudge and TALSA Triplets, as compared to the WAB-R Repetition subscore. The first family of analyses analyzed the relationship among the behavioral variables without any covariates, and included the overall summary measure of aphasia severity, WAB-R AQ. The second family of analyses tested the relationship among the behavioral variables with WAB-R AQ as a covariate, in order to examine the residual relationship among variables when overall language ability was taken into account. We corrected for comparisons using a stringent Bonferroni correction for multiple comparisons, separately within each family of analyses.

We then performed both lesion-symptom mapping analyses (based on proportion damage to each region) and connectome-based lesion-symptom mapping analyses (based on connection strength between each pair of regions) using regression models in NiiStat (https://www.nitrc.org/projects/niistat/). We performed two primary analyses: TALSA Triplets with WAB-R AudWordRec and PALPA RhymeJudge as covariates to control for perceptual processing and overall task performance demands and thus isolate WM load, and Noncanonical sentence comprehension with the residual TALSA Triplets variable as a covariate to control for WM and isolate linguistic processing. Secondary analyses assessing the raw scores for each of these behavioral variables are reported in Supplementary Materials.

All analyses used the full set of 103 participants except for the two analyses involving Noncanonical sentence comprehension, which involved the subset of 78 participants who performed the NAVS Sentence Comprehension task. Lesion-symptom and connectome-symptom mapping analyses were performed on voxels/regions that were damaged in at least 10% of sample in order for maximum spatial reliability. ^74,75^ Connectome-symptom mapping analyses were performed on connections within the left hemisphere only in order to keep the number of independent comparisons within a manageable level.^76^ Lesion-symptom mapping analyses were performed at the voxel level, incorporating overall lesion volume as a covariate to provide strongest spatial inference.^77^ By contrast, all connectome-based analyses did not incorporate lesion volume as a covariate lesion volume, as this was already factored into our analyses by removing damaged tissue from estimated tractographies.^76^ All analyses were corrected for multiple comparisons using permutation testing (4,000 permutations).

## Results

Behavioral results shown in Table 2. Voxel-based lesion-symptom mapping results are shown in Figure 2 and listed in Table 3. For visualization purposes, we display the corrected results along with uncorrected results using a threshold of p < 0.001, in order to provide information about the underlying spatial dimensions of lesion correlates. Significant connectome-based lesion-symptom mapping results showing the relationship between connection strength and behavioral impairments and their associated tractograms are shown in Figure 3.

**Table 2.**
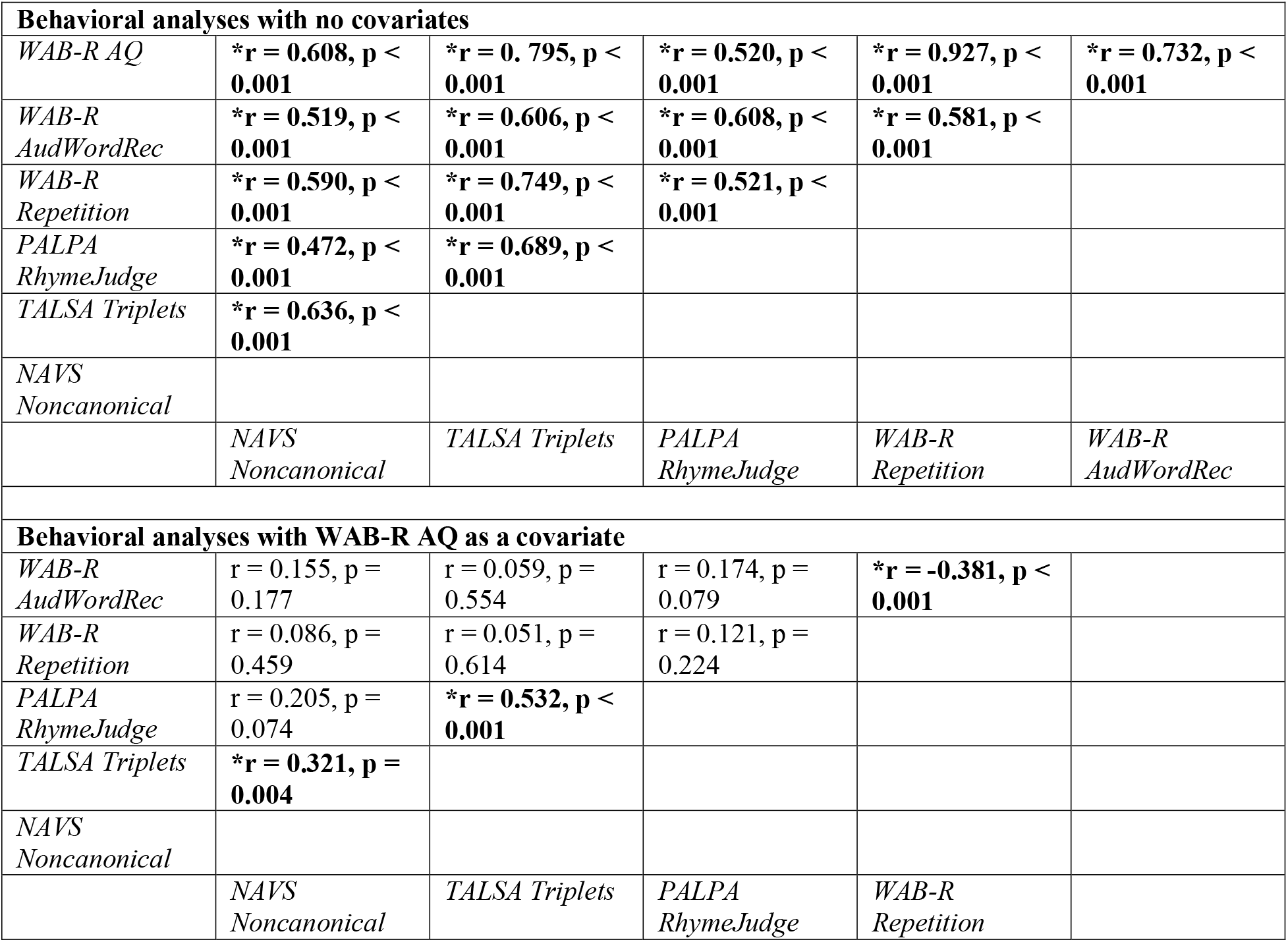
Behavioral correlation results. **\*** and **bold font** indicate significant results. TOP: behavioral correlation analyses, without including any covariates, with an adjusted alpha of p < 0.003 (0.05 adjusted by 15 comparisons) when incorporating a Bonferroni correction for multiple comparisons. BOTTOM: behavioral correlation analyses, incorporating WAB-R AQ as a covariate, with an adjusted alpha of p < 0.005 (0.05 adjusted by 10 comparisons), when incorporating a Bonferroni correction for multiple comparisons.

**Table 3.** Location and size of clusters with at least 20 significant voxels after correction for multiple comparisons using permutation testing (4,000 permutations). Peak MNI coordinates and Peak Z-statistic reflect the location and statistical strength of the single peak voxel within each cluster.

| Region significantly related to behavioral variable | Cluster size (mm<sup>3</sup>) | Peak MNI Coordinates | Peak Z-statistic |
| --- | --- | --- | --- |
| *TALSA Triplets with WAB-R AudWordRec and PALPA RhymeJudge as covariates* |  |  |  |
| White matter directly medial to posterior insula/anterior supramarginal gyrus | 634 | -30, -20, 20 | -5.279 |
| White matter directly medial to postcentral gyrus | 56 | -35, -25, 42 | -4.808 |
| White matter medial to intraparietal sulcus | 37 | -43, -36, 44 | -4.961 |
| Anterior intraparietal sulcus/supramarginal gyrus | 22 | -36, -30, 38 | -4.848 |
| *Noncanonical sentence comprehension with residual Triplets performance (after variance associated WAB-R AudWordRec and PALPA RhymeJudge removed) covaried out* |  |  |  |
| Superior and middle temporal gyrus, extending from the posterior end of the sylvian fissure towards the anterior temporal lobe | 12,643 | -47, -14, -16 | -6.003 |
| Anterior superior temporal gyrus | 80 | -45, 3, -18 | -5.046 |

**Figure 2.**
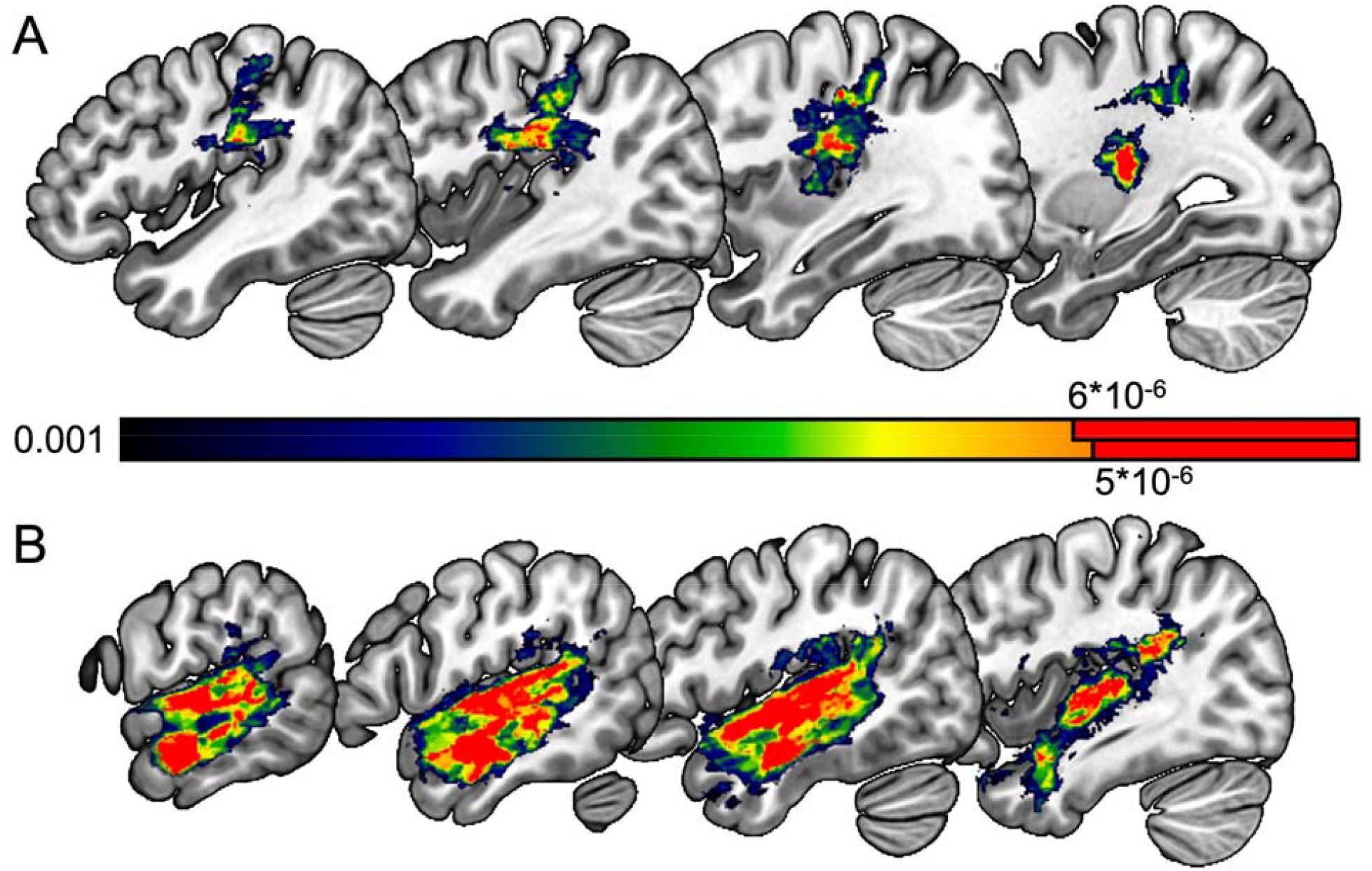
Voxel-wise lesion-symptom mapping results, including total lesion volume as a covariate. A: performance on the TALSA Triplets task, with AudWordRec and RhymeJudge as covariates. B: performance on the Noncanonical sentence comprehension task, with residual Triplets performance as a covariate. Color bar indicates p-value for each voxel. Red color indicates voxels that survived a permutation correction for multiple comparisons (4,000 permutations). Note that the permutation-corrected threshold for the two analyses is different, indicated by the different solid red regions of the color bar.

**Figure 3.**
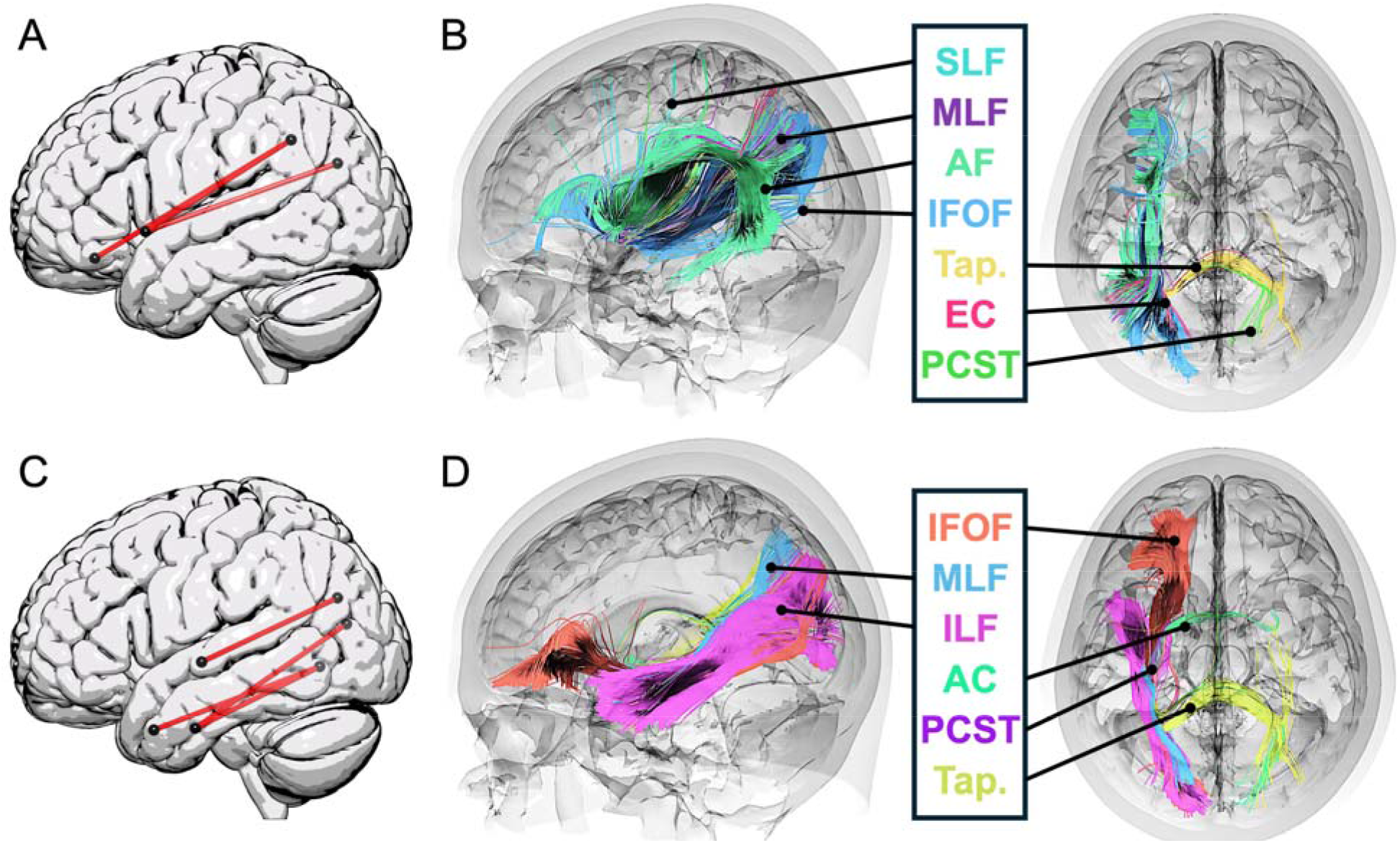
A & C: Significant connectome-based lesion-symptom mapping (CLSM) results for (A) TALSA Triplets with AudWordRec and RhymeJudge covariates, and (C) Noncanonical sentence comprehension with covariate for WM load (residual TALSA Triplets performance). CLSM results are permutation-corrected for multiple comparisons (4,000 permutations). B & D: ROI-based tractography for (B) TALSA Triplets with AudWordRec and RhymeJudge covariates, and (D) Noncanonical sentence comprehension with covariate for WM load (residual TALSA Triplets performance). Fiber-tracking images show all fibers that intersected both pairs of regions implicated in each significant disconnection from CLSM analyses. The specific tract to which each fiber was associated are shown in color-coding, with only tracts containing 20 or more fibers shown. SLF = superior longitudinal fasciculus, AF = arcuate fasciculus, EC = extreme capsule, IFOF = inferior frontal occipital fasciculus, MLF = middle longitudinal fasciculus, ILF = inferior longitudinal fasciculus, AC = anterior commissure, PCST = posterior cortico-striatal tract, Tap. = tapetum.

### Behavioral results

Behavioral correlation analyses without any covariates demonstrated significant and strong relationships among all behavioral variables. We note the particularly strong correlation between overall aphasia severity, WAB-R AQ, and the WAB-R Repetition subscore, which had a Pearson *r* correlation score of 0.927. When including WAB-R AQ as a covariate in partial correlations, most of the correlations were no longer significant. However, the correlation between WAB-R AudWordRec and WAB-R Repetition was reversed, such that there was a significant negative relationship. Interestingly, however, the TALSA Triplets task remained significantly correlated with both the PALPA RhymeJudge and the NAVS Noncanonical sentence comprehension task. While there was possibly still a correlation between PALPA RhymeJudge and Noncanonical sentences, it was no longer significant (p = 0.074). With the inclusion of WAB-R AQ as a covariate, no relationship between Repetition and Noncanonical sentence comprehension was evident.

### Lesion-symptom mapping results

The Triplets task with WAB-R AudWordRec and PALPA RhymeJudge as covariates (Figure 2A) was associated with damage primarily to white matter medial to anterior inferior parietal lobe, straddling the supramarginal gyrus, postcentral gyrus, inferior parietal lobe, and posterior insula. Noncanonical sentence comprehension with residual Triplets performance (after WAB-R AudWordRec and PALPA RhymeJudge were covaried out in multiple regression) as a covariate to remove variance associated with WM load (Figure 2B) was associated with damage throughout the superior and middle temporal gyri, sparing the temporal pole but extending into the posterior inferior parietal lobe.

### Connectome-based lesion symptom mapping results

The Triplets task with behavioral covariates to isolate WM load was significantly associated with three disconnections, each from the broader posterior inferior parietal lobe (angular gyrus/superior occipital gyrus) and more anterior regions, either the anterior insula or lateral frontal-orbital gyrus (Figure 3A) (angular gyrus∼anterior insula, Z = 4.138; angular gyrus∼lateral fronto-orbital gyrus, Z = 4.042; superior occipital gyrus∼anterior insula, Z = 4.215).

Noncanonical sentence comprehension with a covariate for residual Triplets performance measure (after covarying out WAB-R AudWordRec and PALPA RhymeJudge) resulted in four significant disconnections, each between the occipital lobe and central/anterior portions of the inferior and middle temporal gyri (Figure 3C) (superior occipital gyrus∼central superior temporal gyrus, Z = 3.442; cuneus∼anterior inferior temporal gyrus, Z = 3.440; lingual gyrus∼pole of middle temporal gyrus, 3.398; lingual gyrus∼anterior inferior temporal gyrus, Z = 3.394).

To give a more anatomically-realistic rendering of the relevant white matter tracts implicated in verbal WM and syntactic processing, we performed ROI-based fiber tracking in DSI studio https://dsi-studio.labsolver.org/) using the pairs of regions that were significantly disconnected for the TALSA Triplets task with covariates to isolate WM load and the Noncanonical sentence comprehension tasks with the covariate for residual WM load to isolate syntactic processing. We identified streamlines using the tractography included in DSI studio that intersected both regions involved in each disconnection, and aggregated the set of identified streamlines. The results of the ROI-based fiber tracking are shown in Figure 3 (B, D). Impaired performance due to verbal WM load was associated strongly with dorsal white matter tracts such as the arcuate fasciculus and superior longitudinal fasciculus, and to a lesser degree some ventral and subcortical tracts, whereas syntactic comprehension was associated almost entirely with ventral stream tracts such as the inferior frontal occipital fasciculus and inferior longitudinal fasciculus, and to a lesser degree subcortical tracts.

## Discussion

The present study aimed to explore the relationship between the lesion and disconnection correlates of verbal working memory (WM) load and syntactic comprehension ability in people with chronic post-stroke aphasia. While there was a relationship between our rhyme judgment and noncanonical sentence comprehension tasks, both behaviorally and neurobiology, when incorporating controls to isolate the WM load component of the rhyme judgment task and the syntactic component from the noncanonical sentence comprehension task, we found a robust differentiation in the lesion correlates of these abilities. Syntactic comprehension robustly and solely implicated temporal lobe damage and ventral stream disconnection, whereas WM load implicated posterior temporal-inferior parietal damage and dorsal stream disconnection.

### Behavioral Results

There was a significant association between the TALSA Triplets task and both the PALPA RhymeJudge task and the NAVS Noncanonical sentence comprehension task, whether controlling for WAB-R AQ or not. This suggests that the Triplets task taps into aspects of WM load that are distinguishable from overall language impairment. The association between the Triplets task and the NAVS noncanonical sentence comprehension task can be understood based on their shared reliance on verbal WM processes. While the noncanonical sentence comprehension task primarily assesses higher-level linguistic processing, including syntax, it likely engages WM to some extent as well, as indicated by previous research^16,17^. Processing sentences often requires individuals to hold linguistic elements in memory while processing the sentence structure, particularly for noncanonical sentence structures ^2,4,8,13–15,78^, which could involve temporary storage and manipulation of phonological information. Therefore, the significant association between the TALSA Triplets task and the NAVS Noncanonical sentence comprehension task may reflect the shared reliance on WM processes across these tasks. The associations between the TALSA Triplets task and the PALPA Rhyme Judge task are likely driven by the shared cognitive processes of phonological processing. Additionally, the trend towards significance between PALPA RhymeJudge and Noncanonical sentences, despite controlling for WAB-R AQ, hints at shared variance likely driven by verbal WM demands.

There was a robust correlation between the overall severity of aphasia, as measured by the Western Aphasia Battery-Revised Aphasia Quotient (WAB-R AQ), and the WAB-R Repetition subscore. Given the demands of the WAB-R Repetition task, which necessitates proficient auditory processing, phonological processing, motor processing, and short-term memory, this relationship is perhaps unsurprising. It is noteworthy that when accounting for WAB-R AQ, many of the significant behavioral correlations in our data no longer remained significant.

Including WAB-R AQ as a covariate resulted in a significant *negative* correlation between WAB-R AudWordRec and Repetition, which suggests that these variables index at least somewhat meaningfully distinct aspects of language processing. The absence of a significant correlation between repetition and noncanonical sentences after controlling for WAB-R AQ is likely due in part to the fact that the WAB-R Repetition task entails the overt production of real words, involving additional articulatory abilities beyond those necessary for sentence comprehension. It underscores the fact that the WAB-R Repetition subscore might not be a strong index of the WM demands required for comprehending complex sentences.

### Lesion and disconnection mapping results

Lesion-symptom mapping analysis of WM load, assessed through the TALSA Triplets rhyme judgment task with covariates to control for auditory word-picture mapping (AudWordRec) and the ability to perform a metalinguistic rhyme judgment task (PALPA RhymeJudge) yields significant lesion effects in inferior parietal lobe and surrounding cortex. Without these covariates, deficits on the Triplets were not significantly associated with any voxels, underscoring the importance of selecting appropriate covariates to isolate the relevant cognitive-behavioral variable of interest, namely WM load. These results align with prior research implicating these regions in sensory-motor, phonological, and WM-related processes, in functional neuroimaging, neurostimulation, and lesion studies.^79–89^ The association between the inferior parietal lobe and the effect of WM load can be justified by its established role in the phonological loop, a key component of WM responsible for temporarily storing and manipulating auditory information.^19,20^

Prior research has indicated a connection between the inferior frontal cortex and difficulties in performing visual rhyming tasks, which necessitate articulatory rehearsal.^90^ In the realm of Auditory–Verbal Short-Term Memory (STM), the involvement of the frontal cortex has been explored within the framework of the dual stream model.^19,20,83^ According to this model, when individuals perceive speech, it activates sound-based representations in the phonological network. These representations are maintained through corresponding subvocal motor processes in the articulatory network. This continuous cycle of activation and maintenance is facilitated by the sensorimotor interface known as area Spt (Sylvian-parietal-temporal).^81^ Consequently, auditory–verbal STM relies on the dorsal stream for its functioning. Therefore, an intact function of the inferior parietal lobe and surrounding cortex, including area Spt, along with connections to the frontal cortex, facilitates efficient articulatory rehearsal, contributing to the ability to maintain phonological representations in WM. Although we did not identify any lesion correlates for WM load in frontal cortex, suggesting that the inferior parietal lobe is the key processing locus of phonological load, we did find a robust association between disconnection of frontal regions from inferior parietal lobe areas, including prominently dorsal white matter tracts such as the arcuate fasciculus, and impaired behavioral performance. This converges with previous research showing that the frontal cortex is implicated more strongly when more robust manipulation of information is required ^91,92^. This provides provide further support for the overall role of the dorsal stream in WM processes, although the frontal component may be less critical in the phonological storage and maintenance functions of the inferior parietal lobe.

Lesion symptom mapping analyses of Noncanonical sentence comprehension with residual Triplets performance as a covariate implicated superior and middle temporal lobe, spanning from anterior to posterior portions, consistent with previous studies (including data from these same patients) ^46,49,51–53,58,68,93–97^. There were significant disconnections between middle-anterior temporal lobe and posterior temporal lobe and parietal-occipital areas, implicating ventral stream tracts such as the inferior longitudinal fasciculus (ILF) and inferior frontal-occipital fasciculus (IFOF). This suggests that WM and syntactic deficits in aphasia can be meaningfully dissociated by using a similar combination of measures, and that the ventral stream is the primary system for processing receptive syntax as proposed in recent work.^40^ Our results converge with previous work reporting a dissociation between repetition ability and phonological processing (dorsal stream) and language comprehension (ventral stream) using fMRI and diffusion-weighted tractography in healthy adults.^98,99^ While this previous study did not attempt to isolate syntactic processing, it is notable that the dissociation closely mirrors our own using lesion-symptom mapping in people with chronic stroke-based aphasia.

We did not find any regions for which damage was significantly associated with deficits on the PALPA RhymeJudge and WAB-R AudWordRec task when including total lesion volume as a covariate and correcting for multiple comparisons with permutation testing, which contrasts with previous studies that have found significant effects for the same or similar measures.^51,52,76,100–102^ However, some previous studies used additional covariates or did not incorporate a lesion volume covariate. This underscores the importance of considering methodological differences and potential confounding variables when interpreting lesion-symptom mapping results.

### Limitations

Our study involved people with chronic post-stroke aphasia. Thus, the results are limited by the fact that we were only able to examine left hemisphere perisylvian regions with the LSM method. In addition, it is possible that recovery and functional reorganization post-stroke could have affected our results, for example by limiting the implication of frontal regions in our analyses. It is also important to note that linear sequencing demands were low in our rhyme judgment tasks. Thus, these measures tap into the active maintenance of information without necessarily requiring much maintenance of their *order*. It may be the case that memory tasks requiring not just maintenance of stimulus content, but also strict order, would implicate damage to frontal brain systems more directly beyond the disconnection results we obtained.^24,40^ In addition, while conceptual-semantic processing demands in the noncanonical sentence comprehension task were limited, there were clearly still lexical and semantic demands for this task. Future research might seek to test patients on more pure measures of receptive syntax ability, or seek to isolate syntax from semantics more effectively.

### Conclusions

In conclusion, we were able to show a robust dissociation in the lesion and disconnection profile between verbal WM load (posterior temporal-inferior parietal lobe, primarily dorsal stream white matter tracts) and syntactic processing (temporal lobe, primarily ventral stream white matter tracts). The TALSA Triplets Rhyme task with heavy load (3 comparisons), controlled for the overall ability to perform a rhyme judgment task and to identify a visual object given an auditory word input, appears to provide a meaningful operationalization of verbal WM load in people with post-stroke aphasia. As such, it correlates with other measures that involve WM load such as complex, noncanonical sentence comprehension, superior to WAB-R repetition, and can be used to segregate out the verbal WM component of noncanonical sentence comprehension ability from more core aspects of syntactic and semantic processing.

## Data availability

The data that support the findings of this study are available from the corresponding author, upon reasonable request.

## Acknowledgments

We would like to thank Leigh Ann Spell, Allison Croxton, Anna Doyle, Michele Martin, Katie Murphy, and Sara Sayers for their assistance with data collection and scoring.

## Supplementary material

There is no supplementary material associated with this publication.

