## Supplementary Materials for "Verbal working memory and syntactic comprehension segregate into the dorsal and ventral streams"

Deficits in WAB-R Auditory Word Recognition (AudWordRec) (Figure 2, bottom left) were not significantly associated with damage to any regions in our corrected atlas-based analyses. The reduced-threshold, uncorrected voxel-based analysis of WAB-R AudWordRec did reveal weaker effects in the posterior superior temporal sulcus and inferior angular gyrus, suggesting that these effects were merely sub-threshold in the present dataset. Similarly, we did not find any regions that were significantly associated with deficits on the PALPA Auditory Rhyme Judgment (RhymeJudge) task (Figure 2, bottom right) in the analyses corrected for multiple comparisons. However, the uncorrected voxel-based analysis did show some association between deficits on this task and damage to similar regions as the WAB-R AudWordRec task.

By contrast, both the TALSA Triplets task and the NAVS Noncanonical sentence comprehension task produced more robust lesion correlates. Deficits on the Triplets task without any behavioral covariates (Figure 2, top left) were significantly associated with damage to the superior longitudinal fasciculus in the corrected atlas-based analysis, and the reduced threshold voxel-wise analysis revealed additional extent into the inferior parietal lobe. Deficits on Noncanonical sentence comprehension (Figure 2, middle left), were associated with damage to much of the temporal lobe, insula, superior longitudinal fasciculus, supramarginal gyrus and angular gyrus.

The WAB-R AudWordRec task (Figure 3, bottom left), which was not significantly associated with damage to any regions in the corrected analyses, was significantly associated with a large number of disconnections both within the temporal lobe and between the temporal and parietal lobes, also involving subcortical structures. The PALPA RhymeJudge task (Figure 3, bottom right) was significantly associated with two disconnections, one involving two occipital lobe regions bordering on the lateral parietal lobe, and one between posterior middle temporal gyrus and posterior inferior temporal gyrus. The Triplets task on its own was associated with damage to a single region in the lesion-symptom mapping analyses, it was significantly associated with a very large set of perisylvian disconnections (Figure 3, top left). Interestingly, there were no significant disconnections associated with the Noncanonical sentence comprehension task without behavioral covariates.

| Connection | Z |
| --- | --- |
| *WAB-R Auditory Word Recognition (AudWordRec)* | |
| Middle occipital gyrus~thalamus | 3.5599365 |
| Superior occipital gyrus~thalamus | 3.5477300 |
| Superior occipital gyrus~midbrain | 3.4498070 |
| Superior parietal gyrus~cerebral peduncle | 3.4017503 |
| Anterior inferior temporal gyrus~lingual gyrus | 3.3383135 |
| Anterior inferior temporal gyrus~putamen | 3.2654558 |
| Anterior inferior temporal gyrus~cuneus | 3.1738884 |
| Superior occipital gyrus~globus pallidus | 3.1685523 |
| Middle occipital gyrus~midbrain | 3.1670503 |
| Cuneus~globus pallidus | 3.1521325 |
| Cuneus~thalamus | 3.1359823 |
| Cuneus~midbrain | 3.1219935 |
| Pole of middle temporal gyrus~putamen | 3.1044903 |
| Fusiform gyrus~globus pallidus | 3.0854988 |
| Globus pallidus~posterior middle temporal gyrus | 3.0821946 |
| Central superior temporal gyrus~posterior middle temporal gyrus | 3.0708682 |
| *PALPA auditory rhyme judgment (RhymeJudge)* | |
| Superior occipital gyrus~middle occipital gyrus | 3.3609261 |
| Posterior middle temporal gyrus~posterior inferior temporal gyrus | 3.2724634 |
| *TALSA Triplets* | |
| Inferior frontal gyrus pars orbitalis~superior occipital gyrus | 4.6087175 |
| Inferior frontal gyrus pars orbitalis~angular gyrus | 4.1453965 |
| Inferior frontal gyrus pars triangularis~precentral gyrus | 4.1113927 |
| Inferior frontal gyrus pars triangularis~superior occipital gyrus | 4.0984858 |
| Inferior frontal gyrus pars opercularis~angular gyrus | 4.0735854 |
| Inferior frontal gyrus pars triangularis~angular gyrus | 4.0312328 |
| Inferior frontal gyrus pars triangularis~superior parietal gyrus | 3.9800521 |
| Inferior frontal gyrus pars triangularis~middle occipital gyrus | 3.9722085 |
| Inferior frontal gyrus pars orbitalis~postcentral gyrus | 3.9591472 |
| Inferior frontal gyrus pars opercularis~supramarginal gyrus | 3.9330543 |
| Inferior frontal gyrus pars triangularis~supramarginal gyrus | 3.9037185 |
| Inferior frontal gyrus pars opercularis~middle occipital gyrus | 3.9008523 |
| Inferior frontal gyrus pars triangularis~postcentral gyrus | 3.8457129 |
| Anterior superior frontal gyrus~middle occipital gyrus | 3.8405960 |
| Superior parietal gyrus~angular gyrus | 3.8183040 |
| Supramarginal gyrus~angular gyrus | 3.8054954 |
| Lateral fronto-orbital gyrus~hippocampus | 3.8013086 |
| Fusiform gyrus~midbrain | 3.7819662 |
| Lateral fronto-orbital gyrus~superior occipital gyrus | 3.7631529 |
| Precentral gyrus~posterior middle temporal gyrus | 3.7588519 |
| Inferior frontal gyrus pars opercularis~posterior middle temporal gyrus | 3.7435988 |
| Inferior frontal gyrus pars orbitalis~pre-cuneus | 3.7366109 |
| Inferior frontal gyrus pars orbitalis~putamen | 3.7258923 |
| Precentral gyrus~angular gyrus | 3.7039922 |
| Anterior superior frontal gyrus~supramarginal gyrus | 3.6921658 |
| Inferior frontal gyrus pars triangularis~cuneus | 3.6809895 |
| Inferior frontal gyrus pars orbitalis~supramarginal gyrus | 3.6644044 |
| Middle occipital gyrus~thalamus | 3.6465717 |
| Inferior frontal gyrus pars triangularis~posterior superior temporal gyrus | 3.6170475 |
| Anterior superior frontal gyrus~thalamus | 3.6110066 |
| Inferior frontal gyrus pars triangularis~posterior middle temporal gyrus | 3.6076928 |
| Anterior inferior temporal gyrus~thalamus | 3.6027658 |
| Angular gyrus~putamen | 3.5529528 |
| Inferior frontal gyrus pars orbitalis~precentral gyrus | 3.5382141 |
| Inferior frontal gyrus pars orbitalis~superior parietal gyrus | 3.5127060 |
| Middle occipital gyrus~putamen | 3.5008221 |
| Precentral gyrus~supramarginal gyrus | 3.4998049 |
| Hippocampus~midbrain | 3.4732084 |
| Anterior superior frontal gyrus~central middle temporal gyrus | 3.4719221 |
| Globus pallidus~posterior middle temporal gyrus | 3.4695011 |
| Posterior middle frontal gyrus~central middle temporal gyrus | 3.4655704 |
| *Triplets with WAB-R AudWordRec and PALPA RhymeJudge as covariates* | |
| Superior occipital gyrus~anterior insula | 4.2151419 |
| Angular gyrus~anterior insula | 4.1378853 |
| Lateral fronto-orbital gyrus~angular gyrus | 4.0417052 |
| *Noncanonical sentence comprehension* | |
| - | - |
| *Noncanonical sentence comprehension with residual Triplets performance covaried out (after variance associated WAB-R AudWordRec and PALPA RhymeJudge removed, i.e. isolating WM load)* | |
| Central superior temporal gyrus~superior occipital gyrus | 3.4424306 |
| Anterior inferior temporal gyrus~cuneus | 3.4397466 |
| Pole of middle temporal gyrus~lingual gyrus | 3.3979640 |
| Anterior inferior temporal gyrus~lingual gyrus | 3.3941439 |

Supplementary Table X. Disconnection between pairs of regions from the JHU atlas significantly associated with deficits on the behavioral measures of interest after correction for multiple comparisons using permutation testing (4,000 permutations). ~ denotes a bilateral connection between the two indicated regions.

However, the uncorrected voxel-based analyses did find effects in posterior superior temporal sulcus/middle temporal gyrus, posterior superior temporal gyrus, and inferior parietal lobe. This roughly corresponds to the results reported by Pillay et al. for a visual rhyme judgment task without auditory presentation. Interestingly, this convergence for rhyme judgment occurred despite the fact that the tasks used in the present study and by Pillay and colleagues were quite different: our task involved auditory-presented pairs of words, whereas the task in Pillay et al. involved visual text in a design involving the selection of two alternatives that matched a target word. Pillay et al. interpreted these results as supporting a role for phonological processing, not related to WM load, in the cortical territory roughly corresponding to Wernicke’s area in the literature (see ^49,73^. It is possible that more inferior regions of this territory correspond to phonological access ^94,95^, whereas more superior regions (that we identified for the Triplets task, incorporating covariates) correspond to WM load, perhaps invoking motor-articulatory abilities ^73,77,94^.
